# Low-side and multitone suppression in the base of the gerbil cochlea

**DOI:** 10.1101/2024.07.29.605696

**Authors:** C. Elliott Strimbu, Elizabeth S. Olson

## Abstract

The cochlea’s mechanical response to sound stimulation is nonlinear, likely due to saturation of the mechano-electric transduction current that is part of an electromechanical feedback loop. The ability of a second tone or tones to reduce the response to a probe tone is one manifestation of nonlinearity, termed suppression. Using optical coherence tomography to measure motion within the organ of Corti, regional motion variations have been observed. Here, we report on the suppression that occurs within the organ of Corti when a high sound level, low frequency suppressor tone was delivered along with a sweep of discreet single-tones. Responses were measured in the base of the gerbil cochlea at two characteristic frequency locations, with two different directions of observation relative to the sensory tissue’s anatomical axes. Suppression extended over a wide frequency range in the outer hair cell region, whereas it was typically limited to the characteristic frequency peak in the reticular lamina region and at the basilar membrane. Aspects of the observed suppression were consistent with the effect of a saturating nonlinearity. Recent measurements have noted the three-dimensional nature of organ of Corti motion. The effects of suppression observed here could be due to a combination of reduced motion amplitude and altered vibration axis.

**Significance Statement:** The mammalian auditory organ, the cochlea, relies on a nonlinear active process to achieve sensitivity to low-level sounds and sharp frequency selectivity. Recent work using novel interferometric techniques has revealed complex and nonlinear vibration patterns within the cochlea’s sensory tissue. In this study, the motion response to a pure tone was reduced by additional ”suppressor” tones. The observed motion reduction was consistent with the effect of a saturating nonlinearity, possibly compounded by alterations in the axis of cellular vibration, and thus underscoring the 3-dimensional character of cell-based cochlear mechanical activity.

## 1. Introduction

An acoustic stimulus induces a traveling wave in the mammalian auditory organ, the cochlea, which propagates along the cochlear spiral from the base to the apex. Vibration of the sensory tissue, the organ of Corti (OoC) leads to shearing between the apical surface of the OoC, the reticular lamina (RL), and the tectorial membrane (TM), an acellular structure lying atop the OoC. This shearing motion displaces the hair cell stereocilia that protrude from the apical surface of the inner and outer hair cells (IHC and OHC), activating mechanically gated transduction (MET) channels located close to the stereocilias’ tips, converting the displacements into receptor current and potential. Each longitudinal location along the cochlea responds preferentially to a different characteristic frequency (CF) with high frequencies encoded at the base and low frequencies encoded at the apex, known as tonotopic tuning. The cochlea relies on an active process, termed the cochlear amplifier, to boost its mechanical response to low-level sounds. Activity extends the cochlea’s dynamic range over nearly six orders of magnitude in pressure and enhances its frequency selectivity at each location along its tonotopic axis, particularly the high-frequency base. The active process is compressively nonlinear, with that compression likely due to the saturation of the OHC MET current that drives the cochlear amplifier [13]. When two or more tones are delivered, they suppress each other, and two-tone suppression has been documented in basilar membrane (BM) motion and the adjacent pressure, extracellular and intracellular voltage and neural responses *e.g.*, [20, 7, 4, 2, 5]. The suppressive effect of cochlear nonlinearity appears to be important for auditory discrimination *e.g.*, [1]. Cochlear nonlinearity also produces intermodulation distortion products and otoacoustic emissions (DPOAEs) when two or more frequencies are simultaneously presented.

Optical coherence tomography (OCT) and its functional extension, phase sensitive OCT, have advanced the study of cochlear mechanics by allowing simultaneous measurement of vibrations at multiple depths along the OCT optical axis. Traditional interferometic techniques were limited to measurements from the first reflective surface in the light path, typically the BM when recording from the high frequency, basal region of the cochlea. The OCT instrument from Thorlabs is capable of volumetric imaging along with vibrometry, enhancing the ability to identify structures and regions of interest [10, 12, 11]. OCT measurements have revealed differential motions within the OoC, particularly in the region containing the OHCs and Deiters cells, which exhibits sub-CF amplification not observed on the BM [18, 4, 17, 16]. OCT has been used to study two-tone suppression in the mouse cochlea [6], with significant findings regarding the propagation of amplification along the cochlea.

In the experiments presented here, we studied suppression within the OoC by measuring responses to pure tones in the presence of an intense low frequency tone. Responses to zwuis multitone stimuli were recorded at the same locations just after. The intense low frequency suppressor tone is expected to cause significant saturation of MET current, reducing cochlear activity.

Measurements were made near the 25 and 45 kHz CF locations in the base of the gerbil cochlea, at multiple quasi-radial positions across the OoC. Figure 1A shows a schematic image of the OoC in the gerbil base with features that are visible in the OCT B-scan (from the 45 kHz CF location) labelled in Figure 1B. Figures 1C&D show B-scans from the two experimental locations, 25 and 45 kHz CF respectively. The optical (*z* axis) is vertical and the *x* axis was chosen to span the BM approximately radially (left panels of Figs. 1C&D). A volumetric image was rotated to view the *y − z* B-scan (right panels of Figs. 1C&D). This view allows an evaluation of the degree to which the vertical axis is anatomically transverse (perpendicular to BM) versus anatomically longitudinal (parallel to a local segment of the BM spiral). To access the 25 kHz CF location through the round window, the OCT beam was directed deep into the cochlea. The *yz*-plane view of Figure 1C shows that the optical axis for the 25 kHz CF location is predominantly anatomically longitudinal, whereas from the *yz*-plane view in Figure 1D, at the 45 kHz CF location, the optical axis is predominantly anatomically transverse. The program described in [10] quantifies the longitudinal, radial and transverse components of the optical axis, and these numerical *l, r, t* values are noted in the caption.

**Figure 1:**
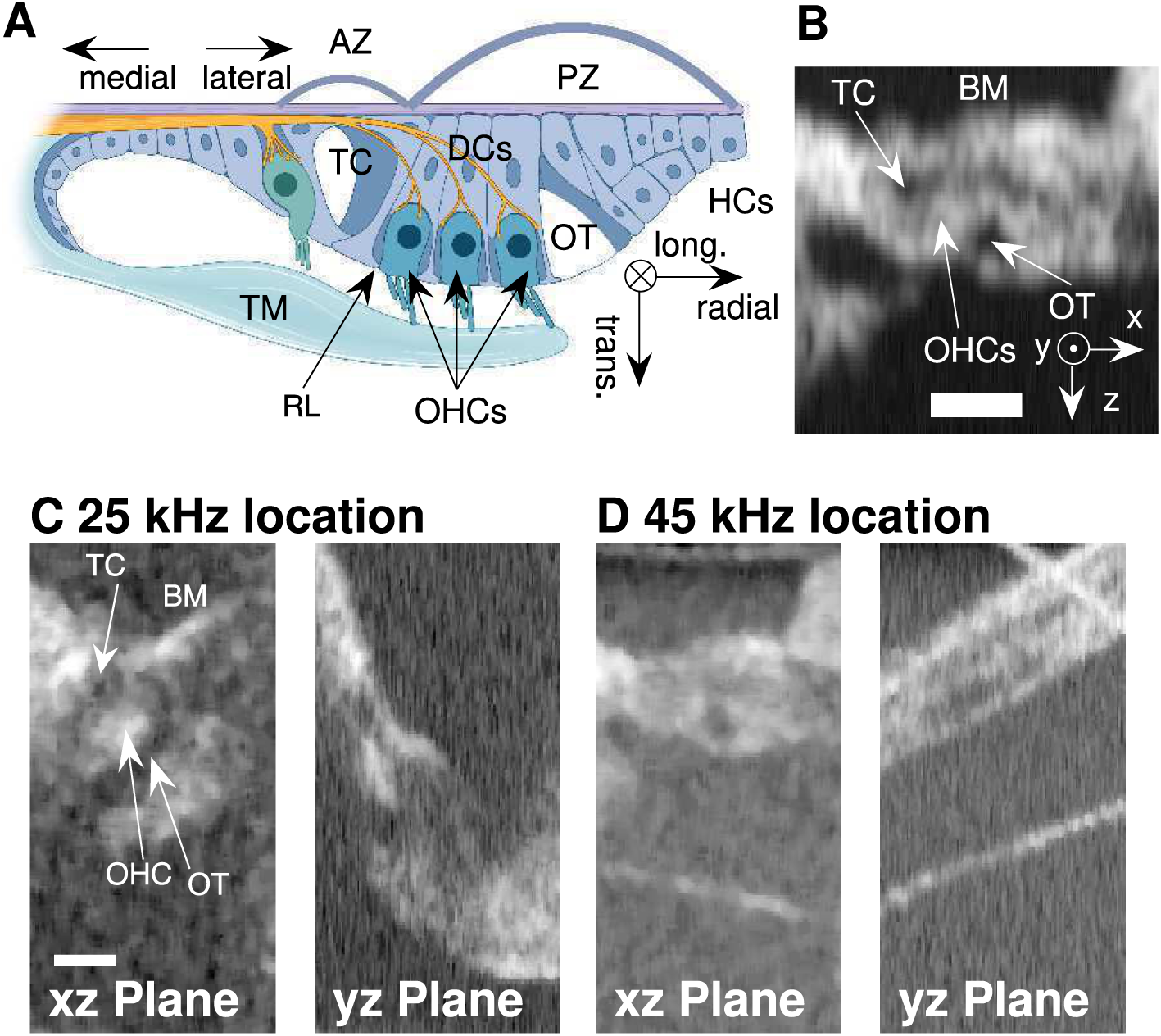
(A): Cartoon cross section of the Organ of Corti with anatomical axes and selected structures labeled: AZ and PZ = arcuate and pectinate zones of the BM, TM = tectorial membrane, TC = tunnel of Corti, OT = outer tunnel, OHCs = outer hair cells, HCs = Hensen’s cells, DCs = Deiters cells, and RL = reticular lamina. The schematic was created with BioRender. In the cochlear base, the fluid-filled spaces within the OoC including the tunnel of Corti and outer tunnel are especially prominent. (B): 2-D B-scan taken at the 45 kHz CF location of the gerbil cochlea. At this location it is possible to align the OCT’s beam giving a nearly transverse view of the OoC; in this specimen the projection of the beam in the transverse direction was 0.98. Scale bar 50 *µ*m. The instrument axes are *z*, along the optical axis, *x* and *y* as indicated in the lower right corner. (C): B-scan from the 25 kHz CF location, with left panel the standard *xz* orientation, and the right panel the *yz* orientation. The inner tunnel, outer tunnel and OHC-region are identified. (D): same as C, for the 45 kHz CF location (see (B) for identifications). (*l, r, t*) values for C are: (0.87,-0.064, 0.49); (*l, r, t*) values for D are: (-0.38, -0.27, 0.89).

Results are presented as tuning curves. DPOAEs were recorded in response to two-tone stimuli and used as a real-time measure of cochlear condition.

## 2. Methods

The experimental protocol was approved by the Columbia University IACUC. Eight adult gerbils of either sex weighing ~ 60 – 80 g were used in this study, and results from six are included in the presented data, three from each CF location. (The suppressor level in the other two was smaller by 10 dB and the suppression was smaller and less illuminating.) The gerbils were anaesthetised with 40 mg/kg ketamine, 40 mg/kg sodium pentobarbitol, and 0.1 mg/kg buprenorphine. Supplemental doses of pentobarbitol were given as needed to maintain areflexia in response to a light hind toe pinch and a second dose of buprenorphine was administered 6 – 8 hours after the start of the surgery. Animals were also given .01 mL of 2% lidocaine at each incision site. The scalp was removed and the skull fixed to a two-axis gonimoeter with dental cement. Animals were tracheotomized to facilitate normal breathing and most were given supplemental oxygen by placing a tube flowing oxygen a few centimeters from the tracheotomy. The left pinna, most of the cartilaginous ear canal, and tissue covering the auditory bulla was resected. The bulla was gently opened after softening the bone with a phosphoric acid gel (Etch-Royale, Pulpdent Corp.) for 5 – 10 minutes. Core body temperature was maintained at 38*^◦^*C with a servo-controlled heating blanket and a rectal thermometer and additional heating was applied to the head with heat lamps and a disposable hand warmer (Hot Hands, Kobayashi Consumer Products) placed on the goniometer. Paper wicks were gently placed within the round window niche to maintain a constant fluid level. Imaging and vibrometry were performed with a ThorLabs Telesto III OCT system equipped with an LSM03 5× objective lens. The ThorImage program was used to position the head and for volumetric (3-dimensional) imaging. The axial resolution of an OCT depends on the central wavelength and bandwidth of the light source and for the Telesto III was reported by the manufacturer to be ~ 4 *µ*m. The lateral resolution is determined by the objective lens and was ~10 *µ*m. For vibrometry, the noise floor is determined by reflectance of the surface of interest and the recording duration, and varied from .05 – 1 nm. The initial conversion of the raw line-camera data to time-locked A-scans (M-scans) was performed with custom software written in C++ using the ThorLabs software development kit; subsequent analyses were performed in custom scripts written in MatLab. Time waveforms were extracted from local maxima in the A-scans [15] and the amplitudes and phases at each frequency were determined by Fourier analysis. For each frequency, the response was deemed significant if its amplitude was three standard deviations above the noise floor measured in ten neighboring frequency bins. The OCT took structural B-scans before and after each set of vibration measurements and we compared the two images to confirm that the preparation was stable, or repeated the measurements if the drift was too severe. Because OCT vibrometry is an interferometric technique, the instrument can only measure vibrations parallel to its light path. In general, the optical axis was not aligned along any of the anatomical (longitudinal, transverse, radial) axes of the cochlea. Frost *et al.* [10] presented a method to determine the mapping between the cochlea’s anatomical axes and the OCT optical axis and we used this method to find the longitudinal, radial, and transverse components of the optical axis for each set of measurements. These are reported in the figure captions as (*l, r, t*) values.

Acoustic stimuli were generated and ear canal pressures were measured using a Tucker Davis Technologies (TDT) system running with a sampling rate of 130 kHz (130.20833 kS/s). Sound was delivered closed-field using a Radio Shack tweeter and the pressure in the ear canal was measured with a Sokolich ultrasonic microphone. The acoustic system was calibrated at the start of each experiment and the calibration was repeated as needed. In each experiment three distinct stimuli were used: pure tones (referred to as a “frequency sweep”), the same pure tones in the presence of a low-side suppressor (referred to as a “sweep + suppressor”), and multitone zwuis complexes [19, 20]. For the sweep and sweep + suppressor measurements, 32 equally spaced frequencies were presented for 62.9 ms each (2^13^ samples) including 1 ms (130 samples) rise/fall times tapered with a cosine-squared envelope for a total recording time of ~2 s. The 3 kHz low-side suppressor was played at 100 dB SPL in six experiments; in two preliminary experiments, one at each CF location, we used a 90 dB suppressor and observed only weak changes in the OoC vibrations. In the zwuis measurements a tone complex containing *N* = 35 sinusoidal components having frequencies of approximately equal spacing (up to a few percent) was presented for ~1 s (2^17^ samples). The frequencies were chosen to have an integer number of cycles in the recording window and to lack harmonics or intermodulation distortion products up to third order. Each sinusoidal component in the tone complex was assigned a random phase, *−π < ϕ_i_ < π* so the total pressure amplitude is *∼* √N, or *∼* 10 log *N* dB SPL, higher than the amplitude at each primary. The SPLs set and noted in the results correspond to the SPL of the individual frequencies. The computer controlling the OCT can process and save a 1 s recording in *∼*10 s and do the same for a 2 s recording in *∼*15 s.

DPOAEs were measured in response to swept two-tone stimuli, *f*_1_ and *f*_2_, held in a fixed ratio of *f*_2_ = 1.2*f*_1_. *f*_2_ ranged from 1 to 48 kHz in 1 kHz steps when recording at the 25 kHz CF location and 2 to 32 kHz in half-octave steps followed by 34 to 60 or 64 kHz in 1 kHz steps when recording at the 45 kHz CF location. The primaries were played at 50 and 70 dB SPL for 64 identical repetitions of 15.7 ms (2048 samples) and averaged in the time domain. Figure 2 shows grouped 2*f*_1_ *f*_2_ DPOAE audiograms. DPOAE levels were generally robust at 50 and 70 dB SPL and stable through the hours of experimentation.

**Figure 2:**
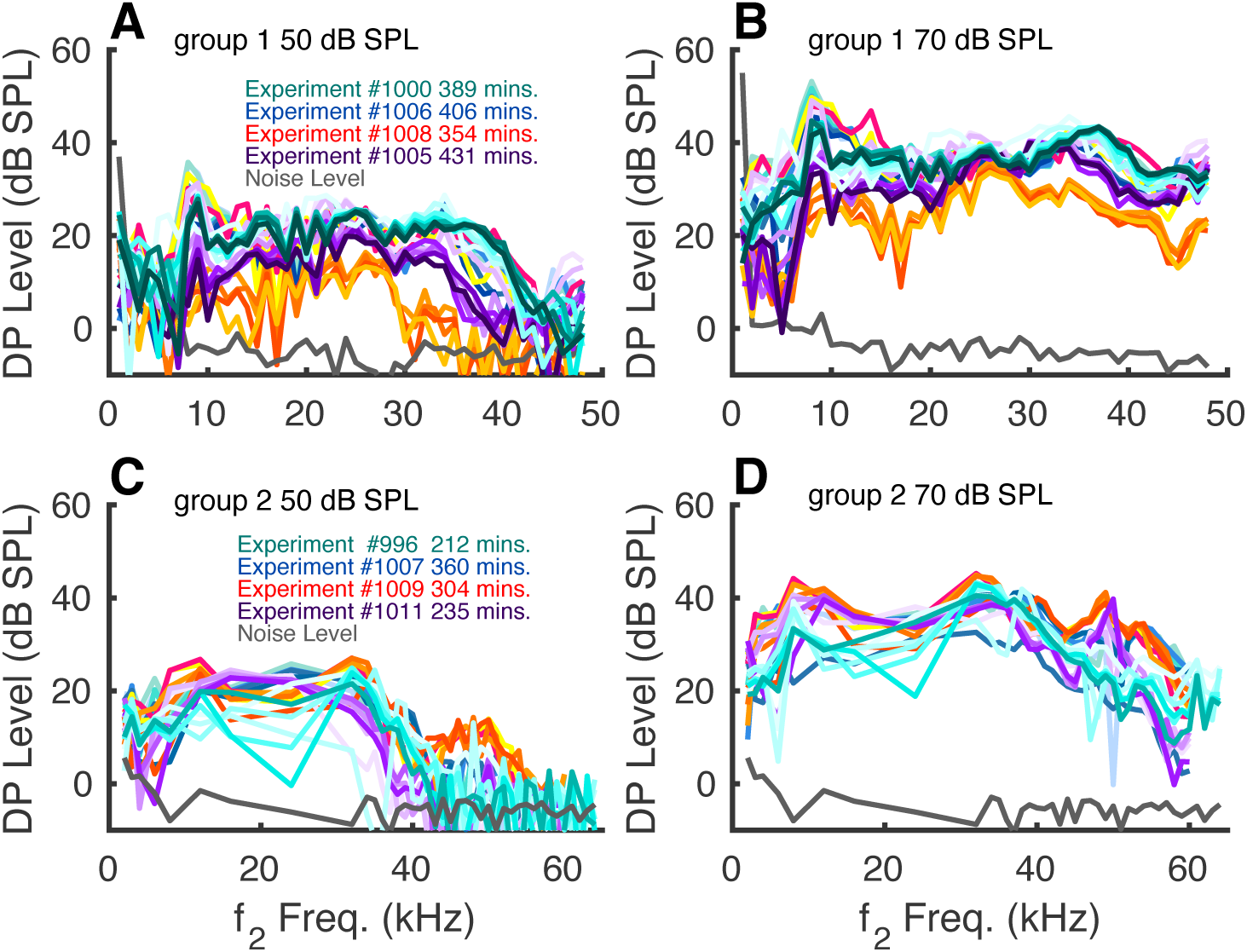
(A – B): DPOAE responses for primary levels of 50 (A) or 70 (B) dB SPL in preparations in which motion was measured at the 25 kHz CF location. Different color schemes are used for each preparation, and trials advancing in time are coded with color saturation; darker = later. The key indicates the latest time. (C – D): Same as above, for preparations in which motion was measured at the 45 kHz CF location.

In each experiment we measured responses at either the ~ 25 kHz or the ~ 45 kHz CF location. Following an initial set of vibration measurements along an optical axis that included the OHC region, we (1) measured DPOAEs, (2) took a volumetric image for orientation, and then at several positions spanning the BM radially we measured: (3) tuning curves in response to a frequency sweep, (4) tuning curves in response to the same sweep in the presence of the low-side suppressor, and (5) tuning curve responses to a zwuis tone complex. A complete set of these five measurements took about 1 hour, and we repeated the set at a few closely spaced longitudinal positions, ~25 microns apart. We employed slightly different radial spanning patterns at the 25 kHz and 45 kHz CF locations. At the 25 kHz CF location, sweep and sweep + suppressor responses were measured across a 120 *µ*m radial span in 20 *µ*m slices (“slice” refers to measurements from one A-scan in the radially-spaced set of A-scan measurements). Responses to the zwuis tones were taken across the radial span in 10 *µ*m slices. For the sweeps, the frequencies ranged from 5 to 40 kHz and were played from 45 to 85 dB SPL in 10 dB steps. The zwuis complex had 35 frequencies covering the same bandwidth and was played at 40, 53, 67, and 80 dB SPL. At the 45 kHz CF location, vibrations were measured in a 144 *µ*m radial span with the sweeps and sweeps + suppressors measured in 11 slices with a 14.4 *µ*m spacing and the frequencies ranging from 5 to 60 kHz and played from 55 to 85 dB SPL. The zwuis responses were measured in 12 *µ*m - spaced slices and played from 50 to 80 dB SPL in 10 dB steps. While the mechanical responses in the 45 kHz CF location are consistent with healthy cochleae showing sharp tuning and compressive growth, we have not been able to consistently measure responses at SPLs below *∼* 50 dB due to signal:noise constraints.

## 3. Results

### 3.1. Multitone versus Single tone

Figure 3 shows single and multitone responses from the ~ 25 kHz CF location. In the B-scan of Figure 3C the BM is identified as the upper surface, but with this very longitudinal view the anatomy of the OoC is otherwise obscure. As in Figure 1C, the two dark areas are loosely identified with the tunnel of Corti and the outer tunnel, which are medial and lateral to the outer hair cells. The large reflective region between them is the “OHC region,” and with the substantially longitudinal view, each A-scan could course through several rows of outer hair cells. A clear difference between the responses to the single-tone and multitone stimulus types is in the sub-CF responses of the OHC region (B&F). In the single-tone responses (B), the sub-CF gain was almost independent of sound level, with a value close to 100 nm/Pa. In the multitone responses (F) sub-CF nonlinearity resulted in lower gains at higher SPL: At 40 dB SPL the sub-CF gain (with sparse data points) was ~ 100 nm/Pa and it fell monotonically as the sound level increased, to a gain of ~ 30 nm/Pa at 80 dB SPL. Another property of the multitone responses from the OHC region in Figure 3F is a relatively steep (compared to single tone) high frequency fall-off at high SPL. Except for the few points affected by this steep fall-off, for both stimulus types and in all frequency regions, there was greater gain in the OHC region than at the BM. The gain at 45 dB SPL with single-tone was slightly higher than the gain at 40 dB SPL with multitone stimulation, which might seem unexpected because 45 dB SPL would produce more saturation than 40 dB SPL, but is reasonable when the quadrature-additive pressure of the multitones is considered. The phases were similar for the two stimulus types, and the phase difference plots in Figure 3D&G show leads of OHC region relative to BM at sub-CF frequencies, consistent with what we have reported previously [18, 17, 16].

**Figure 3:**
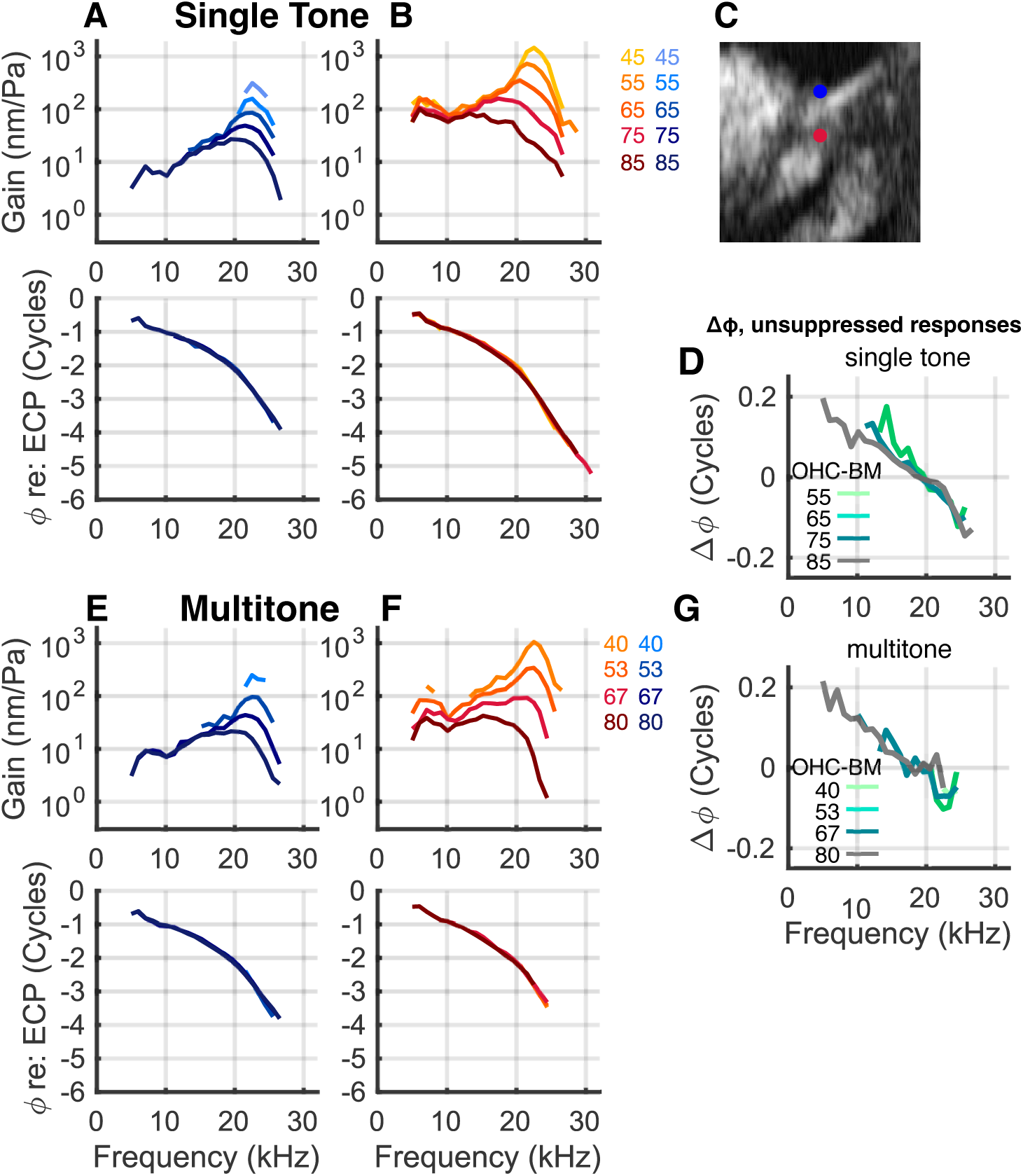
Responses to tones and multitone complexes at the 25 kHz CF location. (C): B-scan. The markers show positions of measurement at the BM (blue) and OHC-region (red). (A) BM and (B) OHC-region tuning curves and phase differences in response to swept tones presented from 45 – 85 dB SPL. (E&F): Responses of the same points to multitone zwuis stimuli presented from 40 – 80 dB SPL. (D&G): Phase differences between OHC and BM responses with single-tone and multitone stimulation respectively. (*l, r, t*) = (0.87, *−*0.064, 0.49). Experiment #1008 runs 20 & 21.

Figure 4 shows single and multitone responses from the ~ 45 kHz CF location. In the B-scan of Figure 4D the viewing axis was nearly transverse and the OoC anatomy is more discernible than in Figure 3. In particular, the surface of the RL can be identified. With the approximately transverse view, the “OHC region” is smaller than with the longitudinal view of Fig. 3D, and each A-scan would generally just pass through at most a single OHC. The responses at both the BM (Fig. 4A&F) and RL region (Fig. 4C&H) were largely similar with the two different stimulus types, and like Figure 3, the greatest response difference between the two stimulus types was in the OHC region (Fig. 4B&G). In the single-tone OHC-region responses (B), the sub-CF gain was almost independent of sound level, at about 40 nm/Pa. In the multitone OHC-region responses (G) sub-CF nonlinearity was apparent and resulted in lower gain at higher SPL. At 50 dB SPL the sub-CF gain was close to 40 nm/Pa and it fell monotonically as the sound level increased, to a gain of 10 nm/Pa at 80 dB SPL. Close to the CF peak, at the lowest SPLs the OHC-region gains were quite similar with the two stimulus types, but at 80 dB SPL the multitone responses show a deep trough that is accompanied by a phase lift (G). Similar but milder behavior is seen in the single-tone OHC-region responses, with a shallow trough observed at 85 dB SPL, accompanied by a ripple in the phase. An explanation for the behavior is in the discussion.

**Figure 4:**
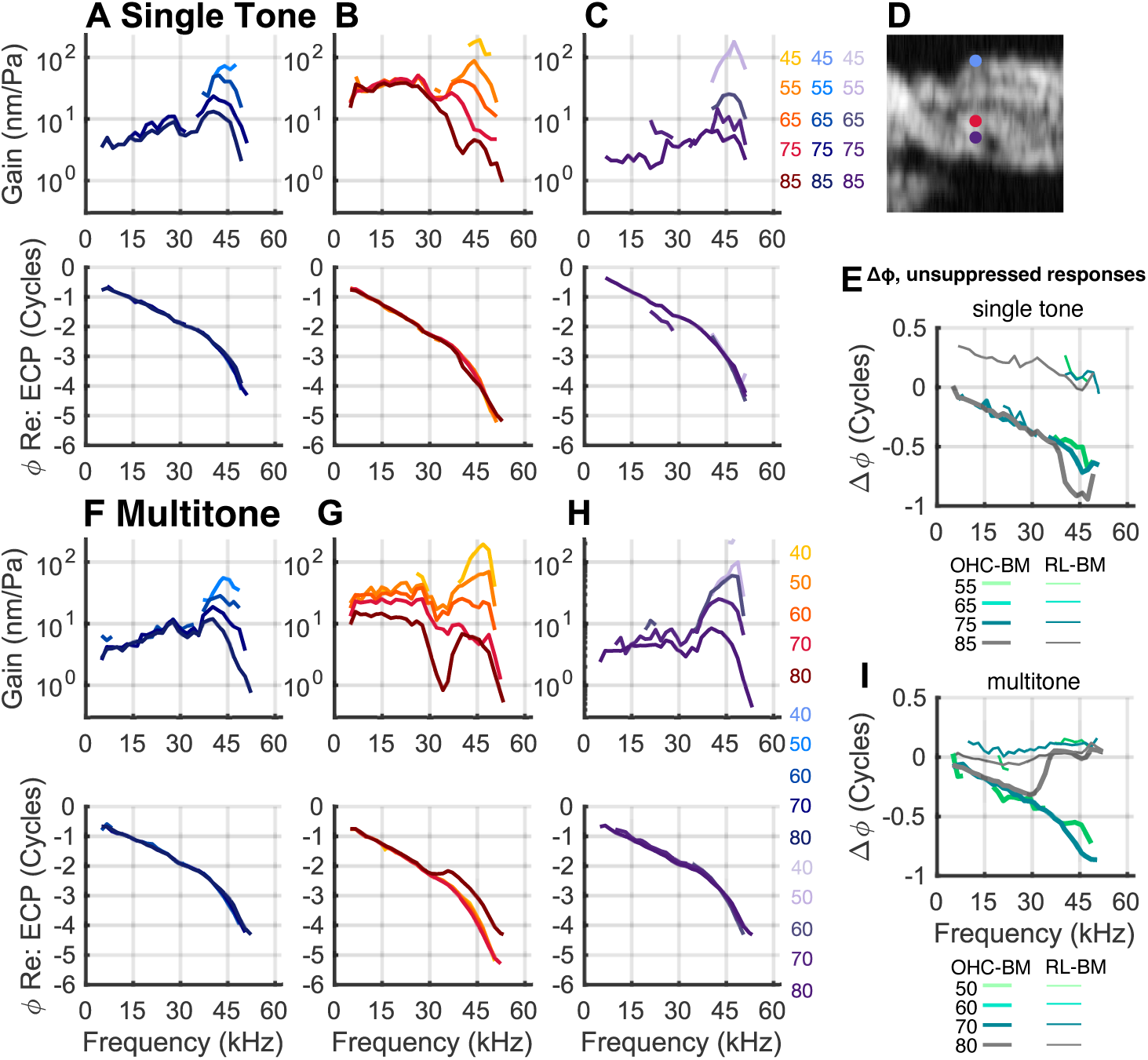
Responses to tones and multitone complexes at the 45 kHz CF location. (D): B-scan. The blue, red and violet markers indicate points on the BM, in the OHC region, and in the RL region respectively. (A – C): The tuning curve gains measured in response to pure tone stimuli at these three positions. (F – H): Tuning curves measured in response to zwuis tone complexes. (E&I) Phase differences between RL-region and BM and OHC-region and BM. (*l, r, t*) = (*−*0.19, *−*0.32, 0.93). Experiment #1009 runs 4 & 6.

An apparent multi/single tone difference between RL responses in Figure 4 is the phase relative to BM phase. With single-tone stimulation, the RL phase led the BM phase (thin lines in Fig. 4E) by increasing amounts as the frequency was reduced from the CF. With multitone stimulation the RL phase was nearly equal to BM phase throughout the frequency range (thin lines in Fig. 4I). However, these different behaviors were not consistently observed and might be due to a small shift in the position of measurement, not the different stimulus types – Cho and Puria [3] observed these two phase behaviors at different radial positions of the RL.

### 3.2. Low side suppressor

The multitone responses in Figures 3&4 form a foundation for the low-side suppressed responses. In the figures to follow we show unsuppressed single-tone responses in reds and the suppressed responses in blues. The suppressor was a 100 dB SPL, 3 kHz tone. In Figures 5&7 we also plot multitone responses along with the low-side-suppressed single-tone responses, to illustrate the similarities and differences between multitone-based suppression and low-side suppression.

**Figure 5:**
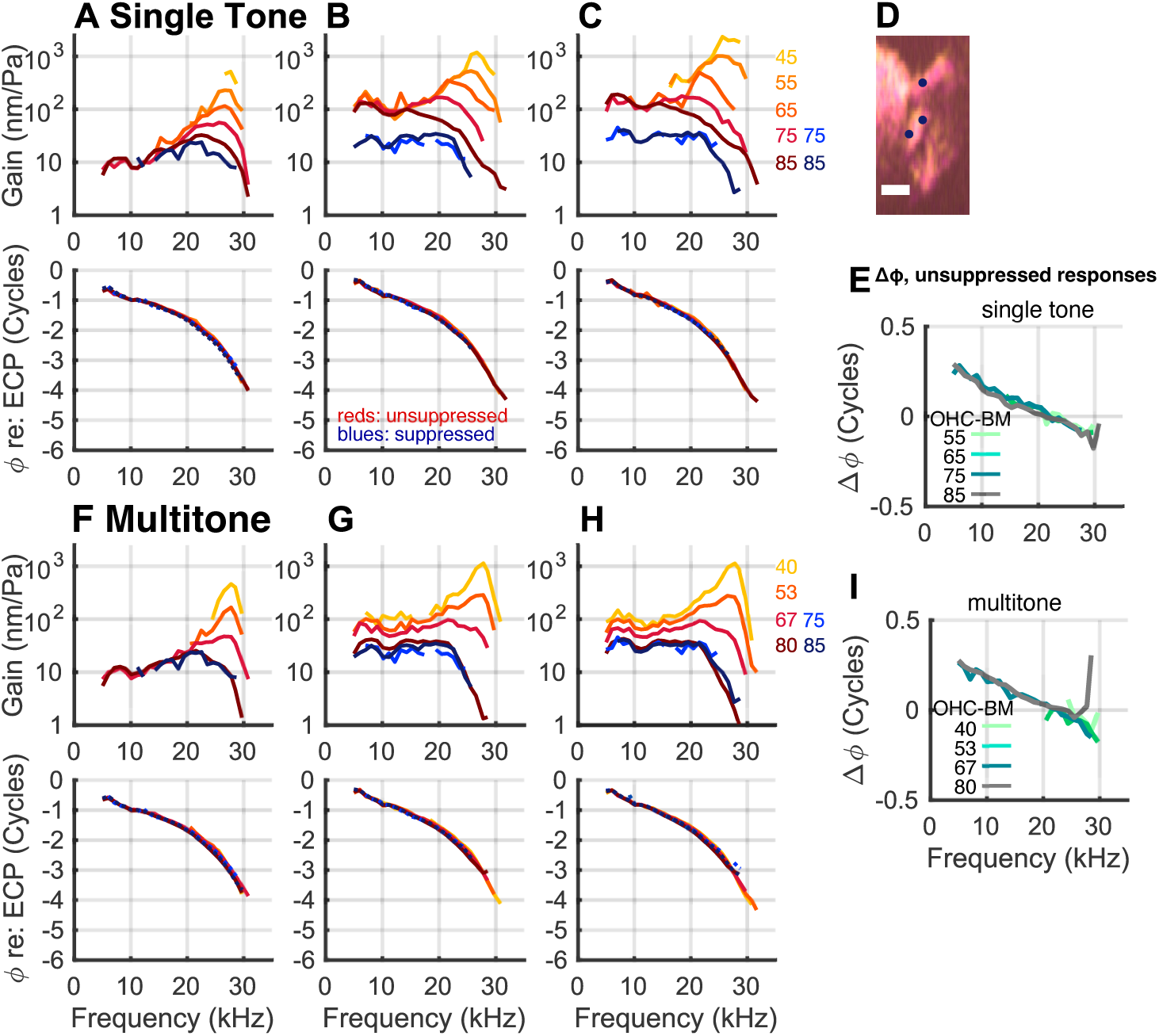
Effects of the low-side suppressor at the 25 kHz CF location. Tuning curves, gain and phase, evoked by single tones at the BM (A) and two locations in the OHC-region (B) and (C). Responses from the single-tones sweep are plotted in reds and the sweep + suppressor are plotted in blues. The points selected are shown in the composite B-scan in D. Scale bar: 40 *µ*m. Responses evoked by the mutitone stimuli are shown in columns (F) – (H) with the suppressed single-tone results overlaid. Experiment #1006, runs 40 (sweep), 41 (sweep+suppressor), and 44 (zwuis). (*l, r, t*) = (0.91, 0.029, 0.41). (E&I) show the phase difference, OHC-region -BM (unsuppressed responses).

Figure 5 shows responses at the 25 kHz CF location. Figure 5A-C shows single-tone responses along with the suppressed responses. Measurement positions were at the BM (A) and within the OHC region (B&C). The B-scan in Figure 5D identifies the positions, A to C, top to bottom. The low-side suppressor reduced the responses substantially, and the suppressed responses were only observable (out of the noise) at the highest stimulus levels, 75 or 85 dB SPL. At the BM (Fig. 5A), the 85 dB SPL suppressed response falls below the 85 dB single-tone level in the CF peak, but sub-CF there is no reduction. In the OHC region (Fig. 5B&C), the low-side-suppressed response is reduced throughout the frequency range, and the 75 and 85 dB SPL responses scaled linearly, so that when displayed as gains they are approximately equal. Even when suppressed, at sub-CF frequencies the OHC-region responses were greater than the BM responses. However, near CF the OHC-region suppressed responses dropped off steeply and became smaller than the BM responses. The response phases, shown in the lower panels, were nearly unaffected by the low-side suppressor; suppressed phase responses are plotted in dotted blue lines, and lie close to the red lines. For completeness, Figure 5E shows the unsuppressed OHC re BM phase, which shows a lead-to-lag that is consistent with previous findings from the 25 kHz CF location (*e.g.*, [16]).

Figure 5F-H shows multitone responses along with suppressed single-tone responses. At both the BM and OHC region, the low-side suppressed responses lie approximately with the 80 dB SPL multitone responses, supporting that the mechanism for suppression, presumably saturation, is similar for multitone and low-side two-tone suppression. Similar to Figure 5E, Figure 5I shows unsuppressed phase differences for the multitone responses, OHC-region re: BM, for the OHC location in (G). The multitone and single-tone phase differences in (E&I) were similiar except for a sudden phase increase in phase at ~ 28 kHz, 80 dB SPL in (I). This increase was due to a phase reversal at 80 dB SPL in the OHC region, which is visible upon close inspection in (G).

Figure 6 shows a more complete set of responses from a different preparation, spanning the OoC radially at the ~ 25 kHz CF location. The B-scan in Figure 6A shows the positions of the reported measurements. It is composed of B-scans from the unsuppressed and suppressed runs plotted together as two different colors and the resulting single color image evinces that there was almost no shifting between runs. Figure 6B shows the A-scans at the various slices, in red for run 20 (single-tones) and blue for run 21 (suppressed single-tones). If there had been no shifting of the cochlea at all between runs the red and blue A-scans would overlie; a small degree of shifting is apparent. The pixel positions noted in Figure 6C-G correspond to the unsuppressed run, and the matching pixels from the suppressed run were close to these but not identical; measurement positions were at local maxima in the A-scans [14, 15]. We show data from the measured points in which both suppressed and unsuppressed results from approximately the same position were detectable (out of the noise).

**Figure 6:**
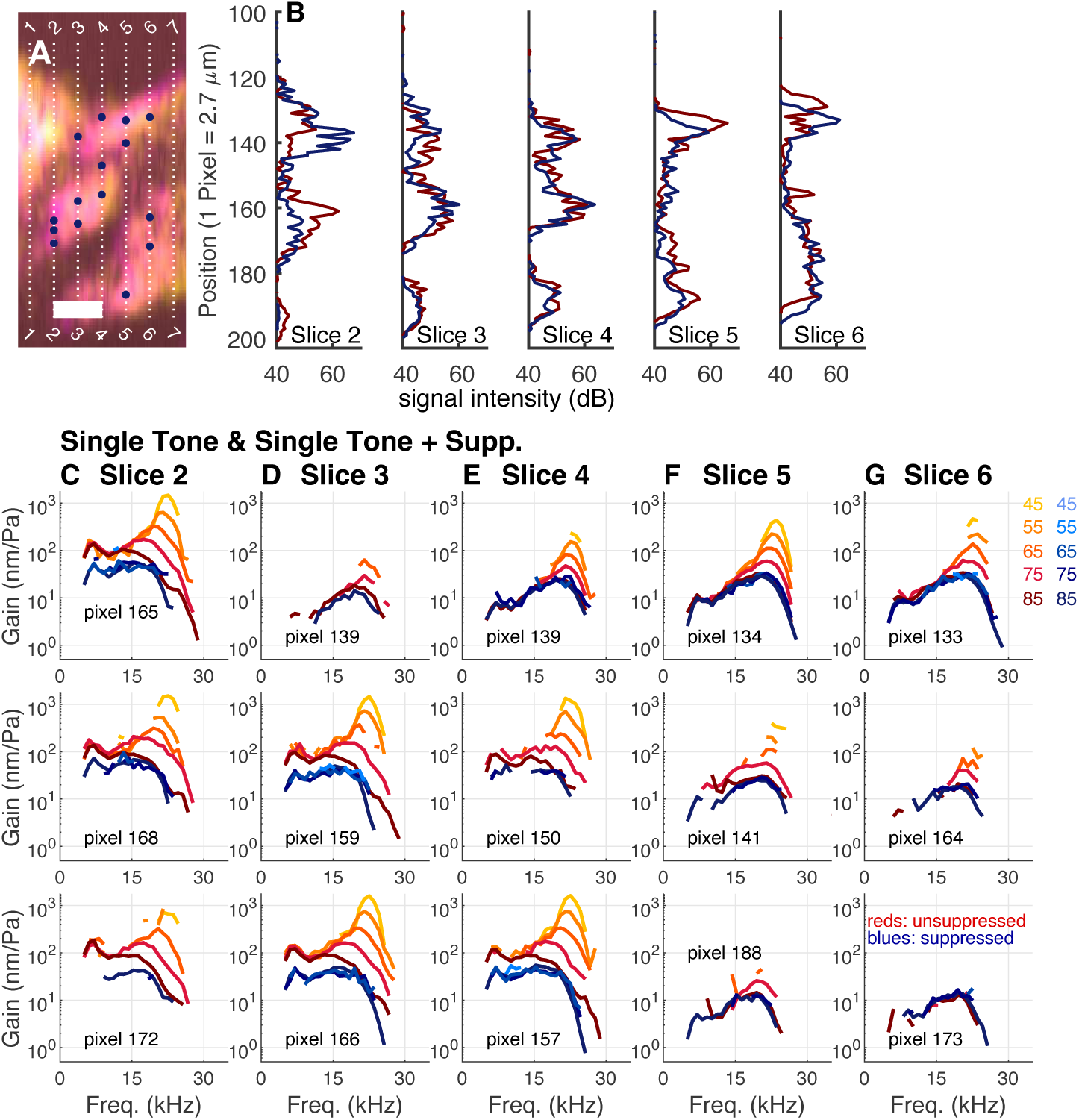
Families of tuning curves measured at multiple positions at the 25 kHz CF location. (A) Composite B-scan taken before the frequency single-tone sweep and before the sweep + suppressor measurement. The blue markers are the points whose gain curves are plotted below. Scale bar 40 *µ*m. Numbered vertical lines indicate slice positions. (B) A-scans for the five slices. A-scans in red correspond to the sweep and those in blue correspond to the sweep + suppressor. Columns (C) – (G) show families of tuning curves evoked by the frequency sweep and sweep + suppressor at different positions along five slices in the quasi-radial direction. As in the previous figure, gains evoked by the sweep are plotted in reds and those evoked by the sweep + suppressor are plotted in blues. Experiment #1008 runs 20 (single-tone sweep) and 21 (sweep+suppressor). (*l, r, t*) = (0.87, *−*0.064, 0.49).

In Figure 6A the large reflective region observed in slices 2, 3 and 4, pixels ~ 150-175, corresponds to the OHC region, and as noted above, with the substantially longitudinal optical axis, each of these slices could have passed through several OHCs. Every measured position from within this region displayed suppressible sub-CF responses. The sub-CF gain of the OHC region was typically ~ 100 nm/Pa unsuppressed, and ~ 60 nm/Pa suppressed. Positions outside the OHC region did not display sub-CF suppression. This included the BM (upper reflective region) in slices 3, 4, 5 and 6, and the lateral region – deep positions in slices 5 and 6. CF-region suppression was observed at all positions, and the suppressed responses scaled linearly – the blue lines showing the suppressed results lie approximately with each other. The unsuppressed CF gain in the OHC region was close to 2000 nm/Pa at 45 dB SPL at all but the deepest OHC region point, slice 2 pixel 172. Suppressed OHC-region responses were low-pass, with gain of at most ~ 60-70 nm/Pa. Close to the CF, at the BM and lateral region, suppressed responses lay nearly with the 85 dB SPL unsuppressed responses. In contrast, close to the CF in the OHC region, suppressed responses dropped more steeply than the 85 dB SPL unsuppressed responses.

Figures 7&8 are correlates of Figures 5&6, from the ~ 45 kHz CF location. At this location the RL region could be identified, and responses from the BM, OHC-region and RL-region are shown. Figure 7A-C shows single-tone responses in reds and suppressed single-tone responses in blues. Figure 7F-H shows multitone responses in reds and suppressed single-tone responses in blues, for comparison. Figure 7A&F correspond to BM, (B&G) to OHC region, (C&H) to RL region, and the B-scan in (D) indicates the positions of measurement. Single-tone suppressed phases are included in (A-C) and (F-G) as dotted blue lines. Figure 7E&I are unsuppressed phase difference results (OHC-BM in thick lines, RL-BM in thin lines) from single-tone and multitone stimuli respectively.

**Figure 7:**
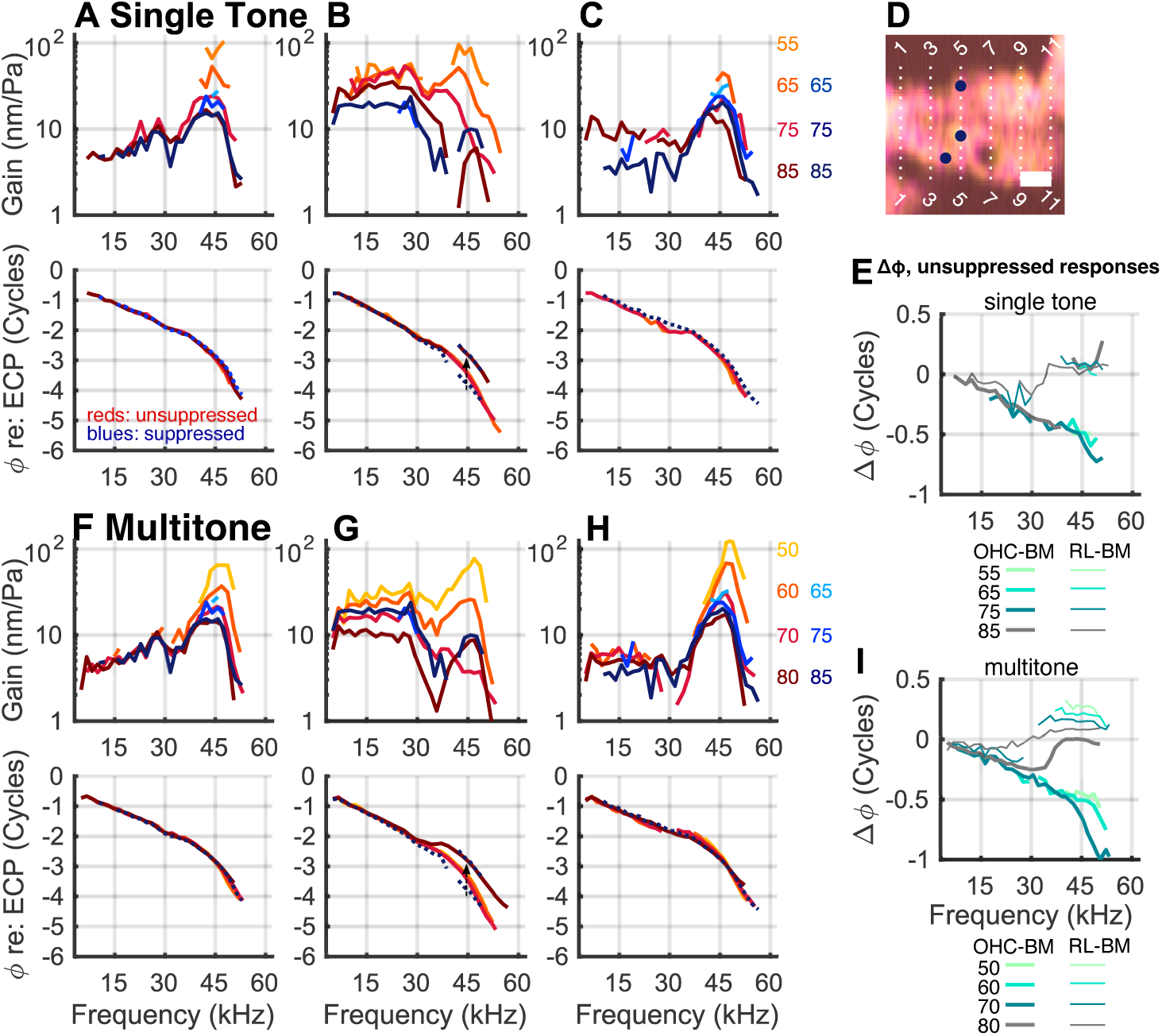
Effects of the low-side suppressor at the 45 kHz CF location. Tuning curves, gain and phase, evoked by pure tones at (A) the BM, (B) OHC/Deiters cell junction, within the OHC-region, and (C) close to the RL. Responses from the single-tones sweep are plotted in reds and the sweep + suppressor are plotted in blues. (F) – (H) show the multitone responses measured at nearby locations with the results of the sweep + suppressor overlaid. Scale bar in D = 28.8 *µ*m. Small blue arrows in the phase plots of (B&G) show a portion of the suppressed curves (blue dotted lines) shifted up a full cycle. (E&I) show the unsuppressed phase differences, OHC-region - BM (thick lines) and RL - BM (thin lines). Experiment #1009, runs 17 (sweeps), 18 (suppressor), and 20 (zwuis). (*l, r, t*) = (*−*0.50, *−*0.47, 0.73).

In both single and multitone responses the BM region showed nonlinearity around CF and approximate linearity sub-CF (Fig. 7A&F). The suppressor reduced the CF peak, but the suppressed peak remained prominent at all SPLs, and was still compressively nonlinear. In the OHC region the sub-CF single-tone responses (B) were nonlinear at the highest SPLs (75, 85 dB) when unsuppressed; when suppressed, the sub-CF responses dropped and were linearized (reporting from a small frequency span around 30 kHz where two SPL responses were observable). In the OHC region the sub-CF multitone responses (G) were nonlinear at all SPLs and the suppressed single-tone responses lay close to the 70 dB SPL multitone response. In the OHC-region CF peak, the single-tone responses (B) were nonlinear when unsuppressed, with a steep fall-off that eliminated the CF peak at 75 dB SPL. However, in both the suppressed and unsuppressed responses, at 85 dB SPL at frequencies above a steep fall-off, a CF peak emerged. These emergent peaks were coincident with a phase lift (lower panel in Fig. 7B). A similar emergent CF peak and phase lift appeared in the multitone OHC gain (G). In Figure 7B&G a portion of the the suppressed phase data is also shown shifted up one full cycle, to show that the suppressed data lay with the lifted phase of the unsuppressed high SPL data. The phase lift is apparent in Figure 7E&I, where the OHC-BM phase (thicker lines) typically decreased from 0 to -0.5 cycles as the CF was approached, but at the highest SPL (thick gray lines), 80 dB for multitone, 85 dB for single tone, abruptly shifted to a phase of ~ 0, beginning at ~ 35 kHz. The significance of this lift was discussed in Strimbu *et al.*, 2024 [16] and will be reviewed in the discussion.

Figure 7C&H show RL-region data. In the single tone gains of (C), sub-CF responses were linear (reporting from a sub-CF frequency span around 25-30 kHz where 75 dB gain lay with the 85 dB gain). The low-side suppressor reduced the sub-CF gain slightly. In the multitone responses the sub-CF gain at the RL (H) was linear and the suppressed single-tone gain lay with the multitone gain. Close to CF in the RL region, the suppressed single-tone gains lay close to the 80 dB SPL multitone gains (H). The RL-region CF peak remained prominent with a degree of compressive nonlinearity even at high SPL and with suppression (both low-side and multitone), similar to the observation at the BM (A&F). The RL-BM phase differences (unsuppressed responses) are the thin lines in Figure 7E&I. The phase differences were slightly negative sub-CF, growing to slightly positive close to CF. The data were more complete for the multitone responses, and these showed a level dependence in phase at frequencies close to CF (I). We don’t consider these SPL-dependent phase variations further here. The RL-BM phase differences were smaller than the OHC-BM phase differences, which increased in absolute value to half-cycle and more close to CF [16]. Figure 8 shows a more complete set of responses spanning the OoC radially in a different preparation, at the ~ 45 kHz CF location. Red curves are unsuppressed single-tone respones, blue are single-tone responses in the presence of the low-side suppressor. Based on the B-scans, positions in slices 4 to 7 contain points that are within the RL and/or OHC regions. The responses in this region can differ qualitatively over distances of just a few micrometers [16] and that appears to be influencing the with/without suppression comparisons at some positions. At many positions in Figure 8, at sub-CF frequencies the suppressed responses lay with the unsuppressed responses, and scaling was linear. This included positions in the BM region in all slices, a position at the edge of the lateral region (pixel 52 of slice 7), and several positions in the RL region in slices 4 and 5. The CF responses at all positions were suppressed, but the CF peak persisted at positions where moderate SPL (65 dB) responses were out of the noise (BM positions slice 8 pixel 32 and 36, RL region slice 4 pixel 61, slice 5 pixel 62.)

**Figure 8:**
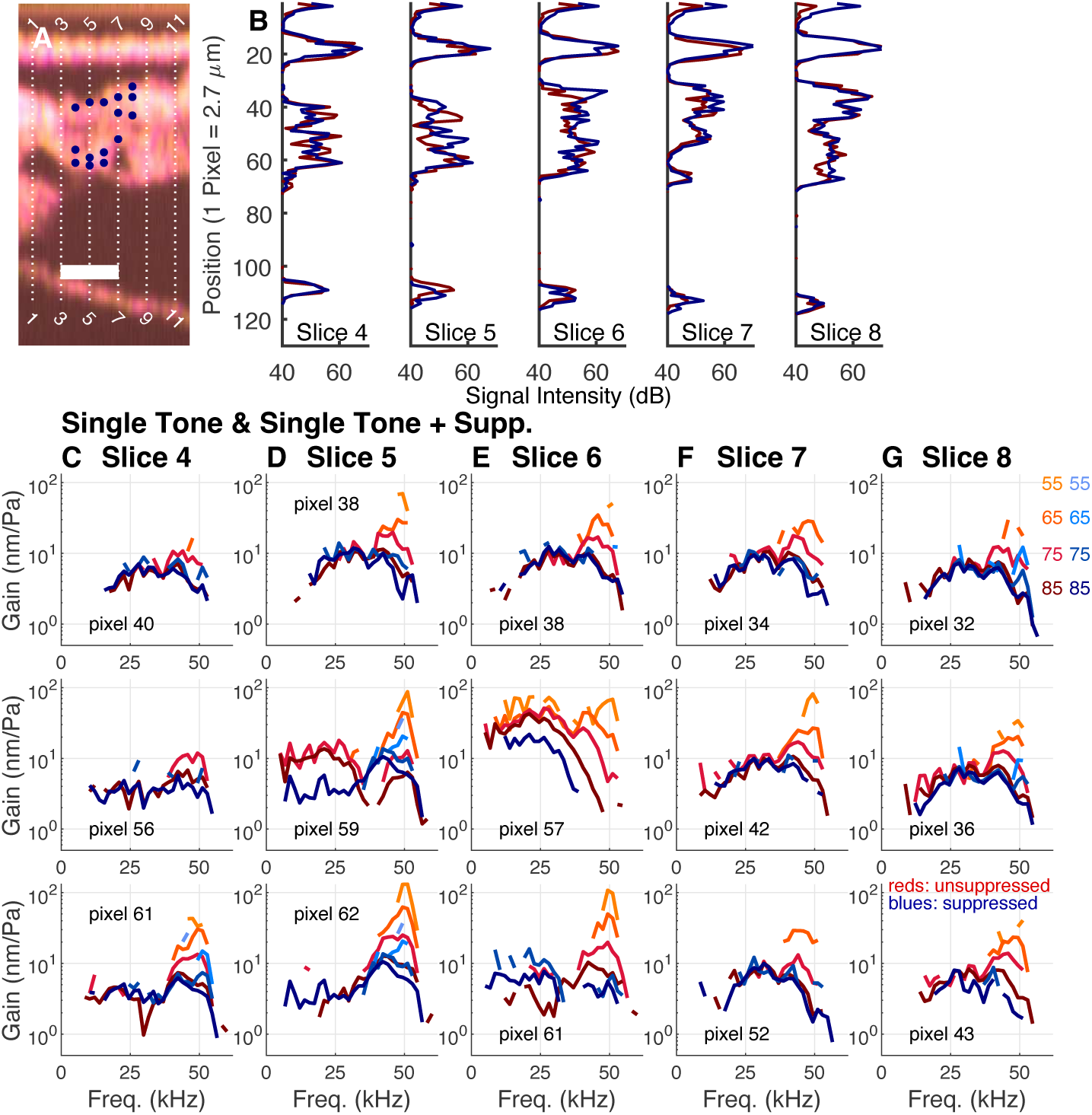
Families of tuning curves measured at different positions at the 45 kHz CF location. The layout of this figure is similar to Figure (6). Experiment 1007, runs 29 (sweeps) and 31 (sweep + suppressor). Scale bar 57.6 *µ*m. (*l, r, t*) = (*−*0.38, *−*0.27, 0.89).

Clear cases of suppressed sub-CF responses in Figure 8 were pixel 59 of slice 5 and pixel 57 of slice 6. Unsuppressed, these data sets showed the elevated sub-CF responses that are characteristic of the OHC region in gerbil. The suppressed gain in slice 6 pixel 57 was low pass and similar in shape to the 85 dB single-tone response, but with a reduced value. The behavior is like that of the suppressed OHC-region responses in Figure 7B, although lacking the emergent CF peak that was observed there. The suppressed response at slice 5 pixel 59 is different – it was not low-pass and the CF peak was still strong at 85 dB SPL – it is similar to the suppressed responses from slice 4 pixel 61 and slice 5 pixel 62, which were positions in the RL region. Slice 5 pixel 59 was close to the border of OHC-body and RL and as noted above, the qualitative behavior can change over small distances in this border region, so this with/without suppression comparison might be influenced by a small position shift (from OHC region to RL region).

### 3.3. Post-mortem Results

The *post mortem* condition renders the cochlea passive, and Figure 9 shows *post mortem* responses from the two CF regions, with (A)–(G) from the 25 kHz and (H)–(P) the 45 kHz CF locations. (A) – (C) show responses to frequency sweeps at the BM and OHC-region with the active vibrations plotted in colors (BM, blues, OHC-region reds) and *post mortem* responses plotted in shades of grey. The phases measured after death are plotted with dotted symbols. (D) – (F) show the responses at roughly the same positions, measured in response to the multitone complexes. The B-scan in (G) shows the measurement positions. *Post mortem* responses (A)–(F) can be compared with the suppressed responses at this CF location in Figure 5. The *post mortem* responses are similar to suppressed responses in being linear and low-pass in the OHC-region, and linear and reduced at the CF, but they are reduced further than they were when suppressed. Thus, suppression had rendered the responses linear, but not passive. The *post mortem* OHC-region responses were smaller than at the BM, likely related to the primarily longitudinal optical axis used at the 25 kHz CF location. The phase responses are similar to active high SPL BM phase responses.

**Figure 9:**
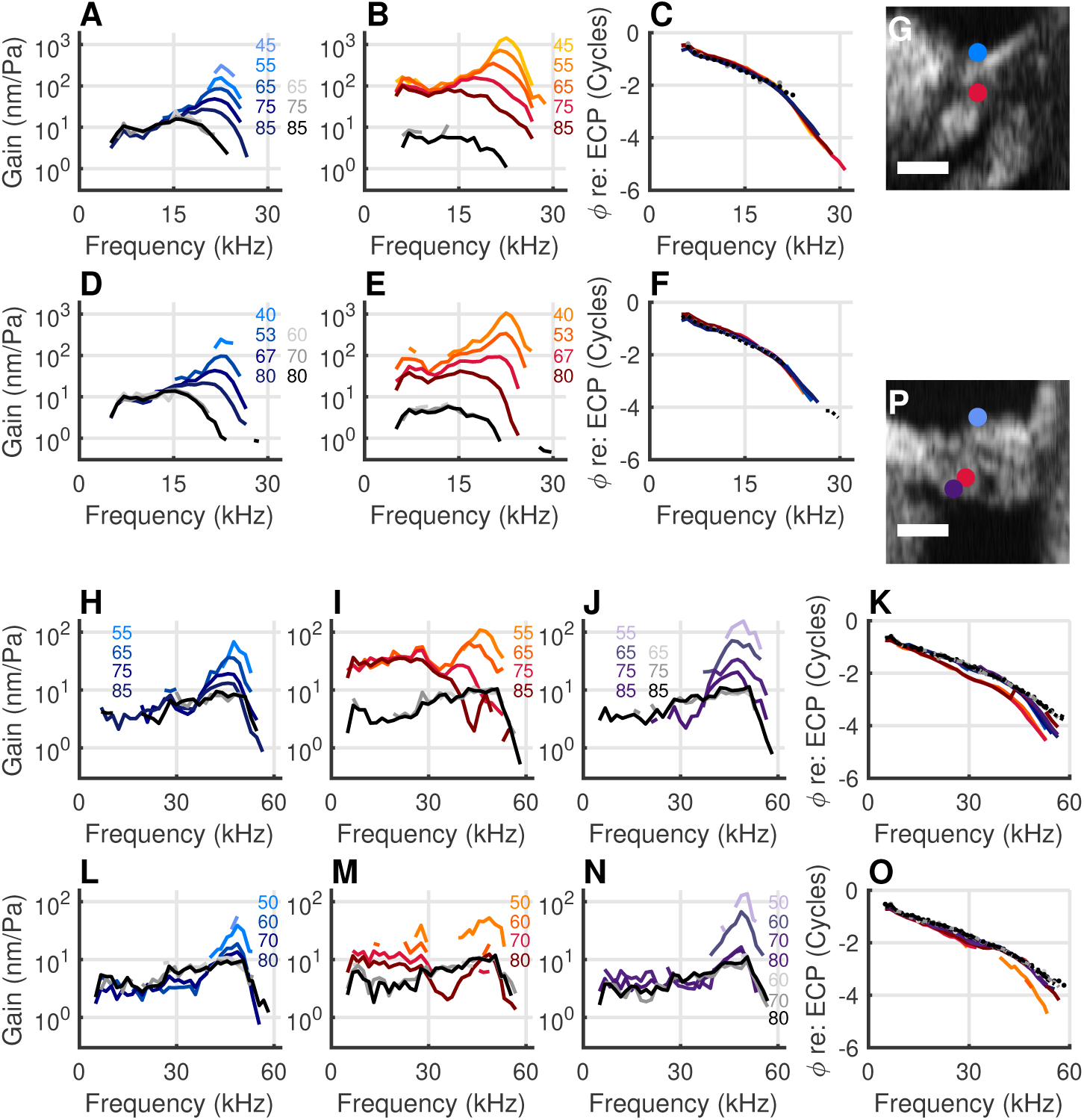
Responses from active and *post mortem* cochleae at the 25 and 45 kHz places in response to the sweeps and zwuis complexes. The top half of the figure shows the responses measured at the 25 kHz place in response to the sweeps and multitone complexes with the BM in blues, the OHC region in reds and the *post mortem* responses plotted in shades of grey. The Bscan in panel (G) shows the two recording locations. Scale bar: 60 *µ*m. Experiment 1008, runs 20 (sweeps), 49 (sweeps *post mortem*), 23 (zwuis), and 51 (zwuis *post mortem*). (*l, r, t*) = (0.87, 0.064, 0.49). The bottom half of the figure shows the responses measured at the 45 kHz place using the same color scheme, with the addition of the RL region in violets. The Bscan in panel (P) shows the three recording locations. Experiment 1009, runs 24 (sweeps), 45 (sweeps *post mortem*), 30 (zwuis), and 48 (zwuis *post mortem*). (*l, r, t*) = (*−*0.45, *−*0.33, 0.82).

Figure 9 (H) – (O) show responses measured at the 45 kHz CF location, with (H) – (K) in response to the sweeps and (L) – (O) to multitone stimuli (BM, blues, OHC-region reds, RL-region violets, *post mortem* grays.) The B-scan in (P) shows the measurement positions. *Post mortem* responses can be compared with the suppressed responses at this CF location in Figure 7. The *post mortem* condition rendered all responses linear. Sub-CF responses were reduced in the OHC region but not changed significantly a the BM or RL. These *post mortem* findings are consistent with those of [3]. The responses still reached a maximum at CF, although the peak response of the *in vivo* condition was eliminated. At this CF location measurements were made with a nearly transverse optical axis, and *post mortem* the responses at the BM, OHC-region and RL-region were all quite similar in size and shape. The *post mortem* phases were largely similar to high-SPL active BM phases, but with a slight lead throughout and smaller phase excursion close to CF.

### 3.4. Grouped Results

Figure 10 plots grouped data of gains at different SPLs at CF and CF/2 for the 25 kHz (A&B respectively) and the 45 kHz CF location (C&D respectively). The three preparations from each CF location went into the grouped data. The two rows in (A&B) are BM and OHC-region; the three rows in (C&D) are BM, OHC-region and RL-region. Unsuppressed single-tone responses are compared to single-tone responses in the presence of the low-side suppressor. Unsuppressed responses are in red symbols, suppressed in blue. The number of data sets included in each symbol is noted above it. If the number of data sets represented in a symbol was less than three, the suppressed/unsuppressed pair was grayed over; the symbols are still useful by providing a gain value, but there were not enough grouped data to reliably use these symbols for a suppressed/unsuppressed comparison. When the number of data sets was three or greater, a statistical comparison was made with a student’s t-test, which tests whether the suppressed and unsuppressed data in a comparison pair were distinct. When the comparison yielded a statistically significant difference, single or double asterisks were placed above the blue and red symbols, depending on the p-value (** for p¡.001, * for p¡.05). Pairs that lack an asterisk were not statistically distinct.

**Figure 10:**
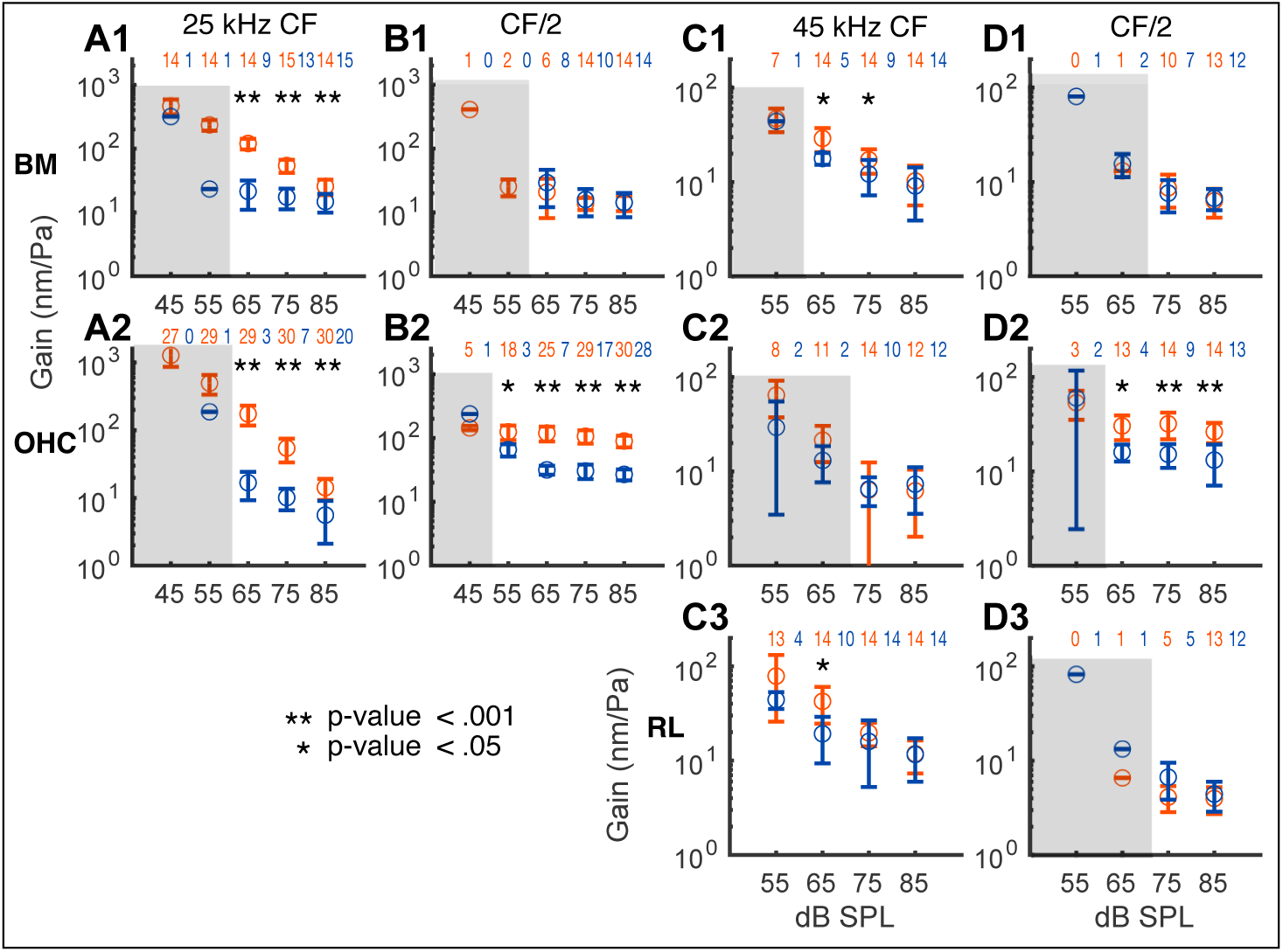
Pooled data from six experiments; three at each tonotopic location. (A&B) Grouped gains at different SPLs at CF and CF/2 for the 25 kHz CF location. (C&D) Same, for the 45 kHz CF location. The two rows in A&B are BM and OHC-region; the three rows in C&D are BM, OHC-region and RL-region respectively. Suppressed responses are in red symbols, unsuppressed in blue. The number of data sets included in each symbol is noted above it. If the number of data sets represented in a symbol was less than three, the suppressed/unsuppressed pair was grayed over due to sparse data. Unsuppressed and suppressed gain values were compared using a Student’s t-test. When the two values were statistically distinct, an asterisk (for p¡.05) or double-asterisk (for p¡.001) was placed above the comparison symbols.

At both the 25 and 45 kHz CF locations, at CF/2 there was no significant suppression at the BM (Fig. 10B1&D1), and there was substantial suppression in the OHC region (B2&D2). In Figure 10B2&D2, the unsuppressed and suppressed gain values were each nearly independent of stimulus level (responses scaled linearly), but the suppressed gain was a smaller value than the unsuppressed gain. At the 25 kHz CF location, at CF both BM and OHC regions showed substantial suppression (A1&A2). At the BM, the CF suppressed response gains were approximately independent of stimulus level, at ~ 20 nm/Pa – the effect of the suppressor was to linearize the responses. At the OHC-region, linearization of suppressed CF responses was also apparent, although less robustly than at the BM. At the 45 kHz CF location both the BM and OHC region showed only a small degree of suppression (C1&C2) – suppression was only significant at the BM. This apparent lack of suppression was likely related to the persistent or emergent CF peak at the 45 kHz location at the highest SPLs, and the fact that there was little measurable suppressed data at SPLs below 75 dB SPL. At the RL-region at the 45 kHz location, CF data were available down to 55 dB SPL and the 55 and 65 dB SPL responses showed suppression (C3). At CF/2 (D3) suppression was not apparent, and gain was approximately equal at 75 and 85 dB SPL (where it could be evaluated), suggesting these responses were not affected by cochlear activity.

## 4. Discussion

We have previously published multitone motion responses from both the 25 and 45 kHz CF locations [18, 17, 16], and the multitone responses here are consistent with our previous findings. In a previous study we compared single and multitone voltage and displacement results from the *∼*25 kHz CF location [9]. The results presented here are consistent with those, but are more complete in frequency span and were taken more quickly due to the sweep strategy employed here, which improved experimental stability.

### 4.1. Expectations from a compressive nonlinearity

In the observations here, the effect of the high-SPL low-side suppressor included linearization of the responses. This effect can be understood within the framework of a saturating input:output curve, as in Figure 11A [8, 7]. Figure 11B shows the output amplitude of a sinewave that is subject to the input:output curve, with the sinewave input amplitude on the *x*-axis. The red curve is the output amplitude of a single sinewave input, the blue curve is the same sinewave’s output amplitude when the input was summed with a second “suppressor” sinewave, of amplitude 10. Two rectangular regions are identified in Figure 11B. The blue rectangle to the left is a region where the red curve scaled linearly, and with the suppressor, it (blue curve) still scaled linearly but at a lower level – ie, with a lower gain. The pink rectangle to the right is a region where the red curve scaled compressively and when suppressed (blue curve), scaled linearly at a reduced level. Both behaviors were observed in our results, and are highlighted in the data of Figure 11C-H, with the left column from the 25 kHz CF location and the right column from the 45 kHz CF location. Each column is from a single run, and these data are shown because some of the suppressed responses were available down to 65 dB SPL – so linearity could be evaluated over the 65-85 dB range of inputs. The blue rectangles identify regions where the unsuppressed responses scaled linearly, and with the suppressor they also scaled linearly, but with reduced gain. The pink rectangles identify regions where the unsuppressed responses scaled nonlinearly and were linearized and reduced in the presence of the suppressor.

**Figure 11:**
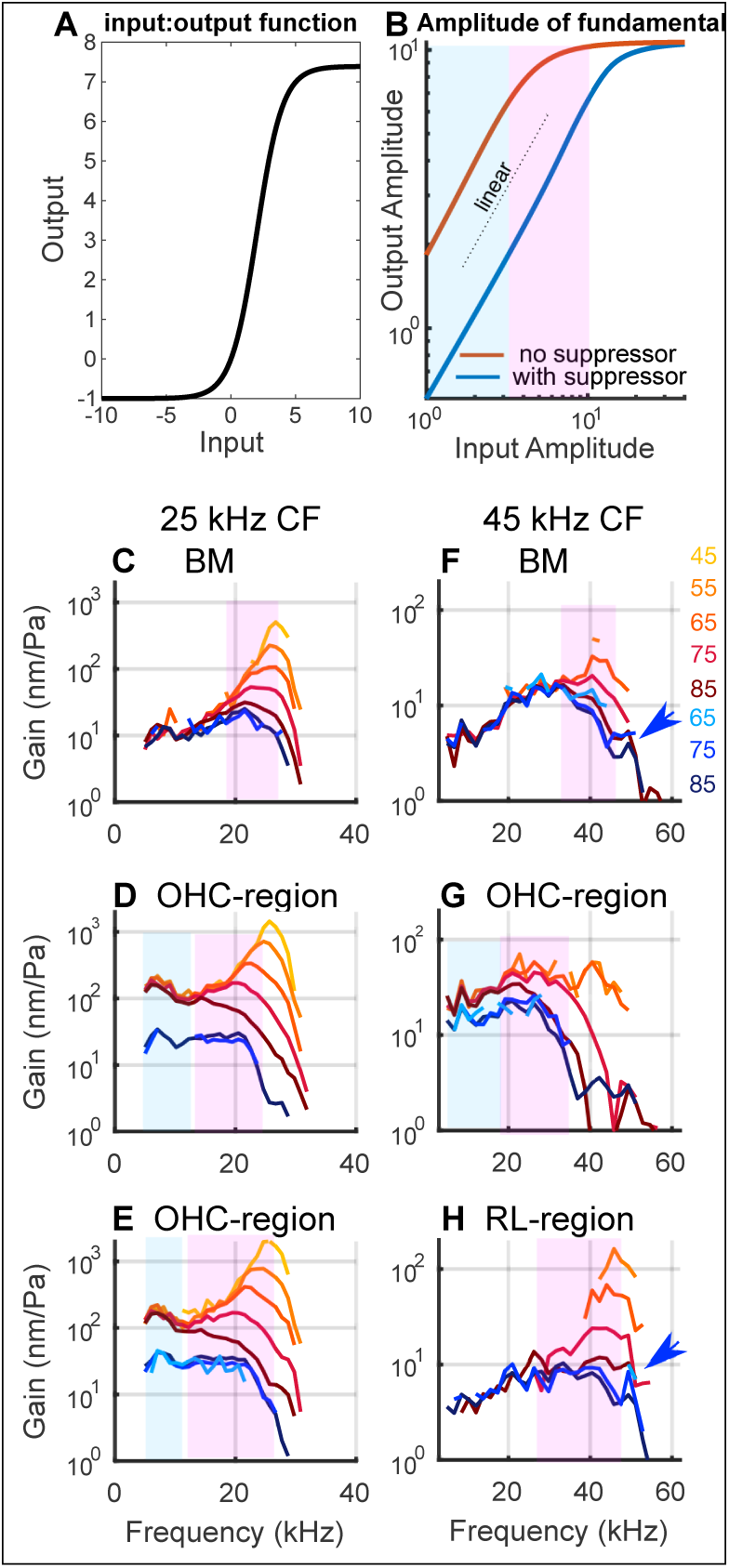
(A) Input:output function of a saturating nonlinearity. (B) Output amplitude of a probe tone of input amplitude as noted on the x-axis. Red = probe tone alone. Blue = probe tone + second (suppressor) tone of amplitude 30. The blue rectangle overlies probe tone responses that were linear in the absence of the suppressor tone, and also linear in the presence of the suppressor tone, but at a lower gain. The pink rectangle overlies probe tone responses that scaled compressively in the absence of the suppressor, and were linearized with reduced gain in the presence of the suppressor tone. (C-E). Single tone and single tone + suppressor responses from the 25 kHz CF location, with reds unsuppressed and blues suppressed responses. The blue and pink rectangles identify the different response characters noted when describing (B). (F-H) Same as (C-E), for the 45 kHz CF location. The blue arrows in (F&H) identify a persistent CF peak in the BM and RL-region responses.

### 4.2. Suppression at CF

The suppressed responses in the OHC-region at frequencies close to CF were remarkable at both the 25 and 45 kHz CF locations. At the 25 kHz CF location, low-side suppression produced a steep fall-off (Fig. 6 slice 3 pixels 159, 166, slice 4 pixel 157). In Figure 5G&H, the high SPL (80 dB) multitone responses show a similar steep fall-off. In previous work [12], measurements along two axes were used to predict overall motion, and showed that, with the optical axis used to probe the 25 kHz CF location, the OHC-region motion could be nearly perpendicular to the optical axis (see also [4, 16]). With that in mind, suppression could modify the overall motion direction, making it even more nearly perpendicular to the viewing axis, and causing the observed steep fall-off. In other words, the size of the observed motion reflects both the motion size and the relationship between the axes of motion and observation, both of which are likely affected by suppression.

At the 45 kHz CF location, the viewing angle was reasonably transverse. Close to CF, OHC-region motion was nearly half a cycle out of phase with BM motion (Fig. 4E&I, Fig. 7E&I). The OHC motion that is measured is a sum of internal active OHC motion and the motion of the BM, the transversely-moving supporting structure of the OoC. At high SPL and/or with low-side suppression, internal OHC motion is reduced, and the motion measured in the OHC-region close to the CF can develop a trough (Fig. 7B&G) which is likely due to the cancellation of internal OHC and BM motion [16]. At the frequencies at and above those of the trough, the phase lifted to join the BM phase (Fig. 7 E&I), indicating that BM motion dominated internal OHC motion. At frequencies above the trough a peak emerged in the OHC-region, which is likely due to the dominance of BM motion.

The blue arrows in Figure 11H identified a persistent nonlinear CF peak that was observed at the BM and RL-region even at high SPL and in the presence of the low-side suppressor. The persistent peak was smaller at the BM than in the RL-region, but still present (see also Figure 8 from the same preparation). It was more prominent in the Figure 7 results from a different preparation and was observed in previous results by us and others at the 45 kHz CF location [16, 3]. (The OHC-region can also display a persistent peak at the 45 kHz CF location; it was not apparent in the suppressed responses in Figure 11, but was in Figure 7.) The 45 kHz CF location was generally less affected by the suppressor than the 25 kHz CF location. For example in Figure 7B&G the suppressor reduced the OHC region gain to the level of the 70 dB SPL multitone gain. In contrast, at the 25 kHz CF location (Fig. 5A&G) the suppressed OHC-region gain was reduced further, to the level of the 80 dB SPL multitone gain. The diminished effect of suppression at the very basal location might be related to this location being so close to the cochlea’s input – sandwiched between the stapes and the round window. Suppression affects the traveling wave as it travels to its CF location [6, 5], and in the very basal location there is little longitudinal space to travel. In a related vein, the *post mortem* responses at the 45 kHz CF location retained a ~ 45 kHz maximum, whereas at the 25 kHz CF location, the BM maximum was shifted downward by about a half octave, and the OHC region responses lacked a peak and became low-pass.

Taken all together, the different suppressed (and *post mortem*) behaviors at the two CF locations are likely attributable to both the longitudinal location being either at or apical of the extreme base, and the optical axis, which was predominantly longitudinal to access the 25 kHz CF location, and predominantly transverse at the 45 kHz CF location.

## 5. Conclusions

Two-tone suppression has long been used to study cochlear activity [13, 7, 20, 6] and OCT allows for the exploration of the effect of suppression on internal OoC components. Our findings regarding BM suppression were like those of previous studies, with suppression affecting the CF region but not the sub-CF region. The OHC-region results showed reduced gain and linearization both at CF and sub-CF, findings that are consistent with previous low-side suppression findings in measurements of extracellular voltage [7]. The RL-region and lateral region of the OoC showed suppressive effects that were qualitatively similar to the BM, with CF-region gain reduction, and little effect of suppression sub-CF. Overall, the effects of the low-frequency high-level suppressor tone were in keeping with the mathematical expectations of a saturating nonlinearity. Certain features of the results, the very steep fall-off of the OHC-region at the 25 kHz CF location, and the persistent peak of the 45 kHz CF location, are likely due to specific cochlear mechanisms, such as the axis of intra-OoC motion, and traveling wave mechanics.

## 6. Author Contributions

ESO and CES designed the experiments. CES conducted the OCT measurements and performed the initial analysis on the data. Both authors authors contributed to further analysis and writing the manuscript.

## Acknowledgements

This work was supported by the NIDCD and the Emil Capita Foundation.

## 7. Declaration of Interests

The authors declare no competing interests.

## Notes

### Competing Interest Statement

The authors have declared no competing interest.

### Summary of Updates

Corrected a typo in the second author's name.

